# Open datasets and code for multi-scale relations on structure, function and neuro-genetics in the human brain

**DOI:** 10.1101/2023.08.04.551953

**Authors:** Antonio Jimenez-Marin, Ibai Diez, Asier Erramuzpe, Sebastiano Stramaglia, Paolo Bonifazi, Jesus M Cortes

## Abstract

The human brain is an extremely complex network of structural and functional connections that operate at multiple spatial and temporal scales. Investigating the relationship between these multi-scale connections is critical to advancing our comprehension of brain function and disorders. However, accurately predicting structural connectivity from its functional counterpart remains a challenging pursuit. One of the major impediments is the lack of public repositories that integrate structural and functional networks at diverse resolutions. Addressing this issue, we provide open datasets and code enabling the examination of the correspondence between structural and functional connectivities at different scales. We present a module-level strategy that overcomes region-level approaches in understanding the structure-function correspondence. Moreover, we also provide additional resources focused on neuro-genetic associations of module-level network metrics, which present promising opportunities to further advance research in the field of network neuroscience, particularly concerning brain disorders.

## Introduction

The precise relationship between brain structural and functional connectivities has puzzled leading researchers in network neuroscience^1–16^. Despite numerous efforts, an accurate prediction of the structural connectivity from its functional counterpart remains still a distant goal. Beyond link-level network comparisons between the two modalities, a modular-level comparison has shown bi-directional advantages in the understanding of brain structure and function^17–29^. Although non-invasive magnetic resonance imaging (MRI) techniques have a limitation in the network resolution that they can resolve^30^, arriving at about one millimeter resolution, multi-scale approaches between brain structural and functional modules have been studied to assess organizational aspects in healthy population^13,26,31–33^, in relation to cognition^34,35^, during development^36^ and aging^37^, or in some pathological conditions^38,39^.

When evaluating the module-level (a.k.a. aggregated-level) correspondence between structure and function, there are two differentiated strategies in relation to the chosen network resolution. One is the optimal strategy, in which a fixed level of network representation is established, for example, by maximizing a certain network metric^40–45^. In an alternative approach, the multi-scale strategy involves employing a nested hierarchy of network resolutions as potential features or input variables to be synergistically integrated for the purpose of addressing a specific problem^35,37,39,46^.

While progress has been made in assessing the module-level correspondence between structural and functional connectivities in the human brain, many questions remain unanswered. Among these inquiries, it is important to assess the extent to which the different modules within structural connectivity favor the emergence of functional modules. Moreover, it is crucial to investigate the presence of dynamic processes that influence the relationship between structural and functional modules. Identifying the scale at which the correspondence between modules in both representations reaches a maximum is also of great relevance. Understanding the insights of the pathological brain and the extent to which the interplay between structure and function is altered within this context are pressing matters. Additionally, elucidating the underlying mechanism responsible for such alterations is crucial. However, a major limitation for precise answers is the lack of public repositories that combine structural and functional networks at different levels of resolution, together with useful and practical code for studying the module-level relation between the two network modalities.

This study aims to address this need by providing datasets and code to the scientific community, enabling progress in addressing these questions, which constitute a central focus of research in our laboratory. Specifically, in this article, we initially introduce a modeling approach to investigate the interplay between structure and function by parameterizing the degree of overlap between the two classes of connectivity. Secondly, building upon the methodology described in^24^, we extend it to examine the module-level correspondence between the two connectivity matrices. Finally, we have extended the study on the relationship between brain structure and function by sharing also neuro-genetic code and data, delving into the underlying biological and molecular foundations of brain-related disorders, similarly to previous work^47–57^.

## Results

Structural and functional connectivity matrices at various resolutions were built making use of brain images from the open dataset “Max Planck Institut Leipzig Mind-Brain-Body Dataset” – LEMON^58^, well-known for having high-quality multimodal acquisitions and preprocessed MRI sequences. Rather than using different subjects across lifespan, we specifically selected 136 young participants, aged between 20 and 30 years old, who had both the preprocessed rs-fMRI sequence and DWI sequence available. Next, we modeled the amount of interplay between structure and function connectivities using a single fusion parameter, *γ*. When *γ* equals 0 or 1, we have purely structural or functional connectivity, respectively. In intermediate situations, the amount of overlapping connectivity is modulated by *γ*.

We next processed the raw images following standard neuroimaging pipelines to obtain structural connectivity (SC) and functional connectivity (FC) matrices (Figure 1A). Subsequently, we built population connectivity matrices by choosing, for each link in the matrix, the median value across all the equivalent links in the individual connectivity matrices. Furthermore, and similar to the methodology described in^24^, for each value of the fusion parameter *γ*, we applied hierarchical agglomerative clustering to the resulting *γ*-fused structure-function matrix, *γ*SFC (Figure 1B).

**Figure 1.**
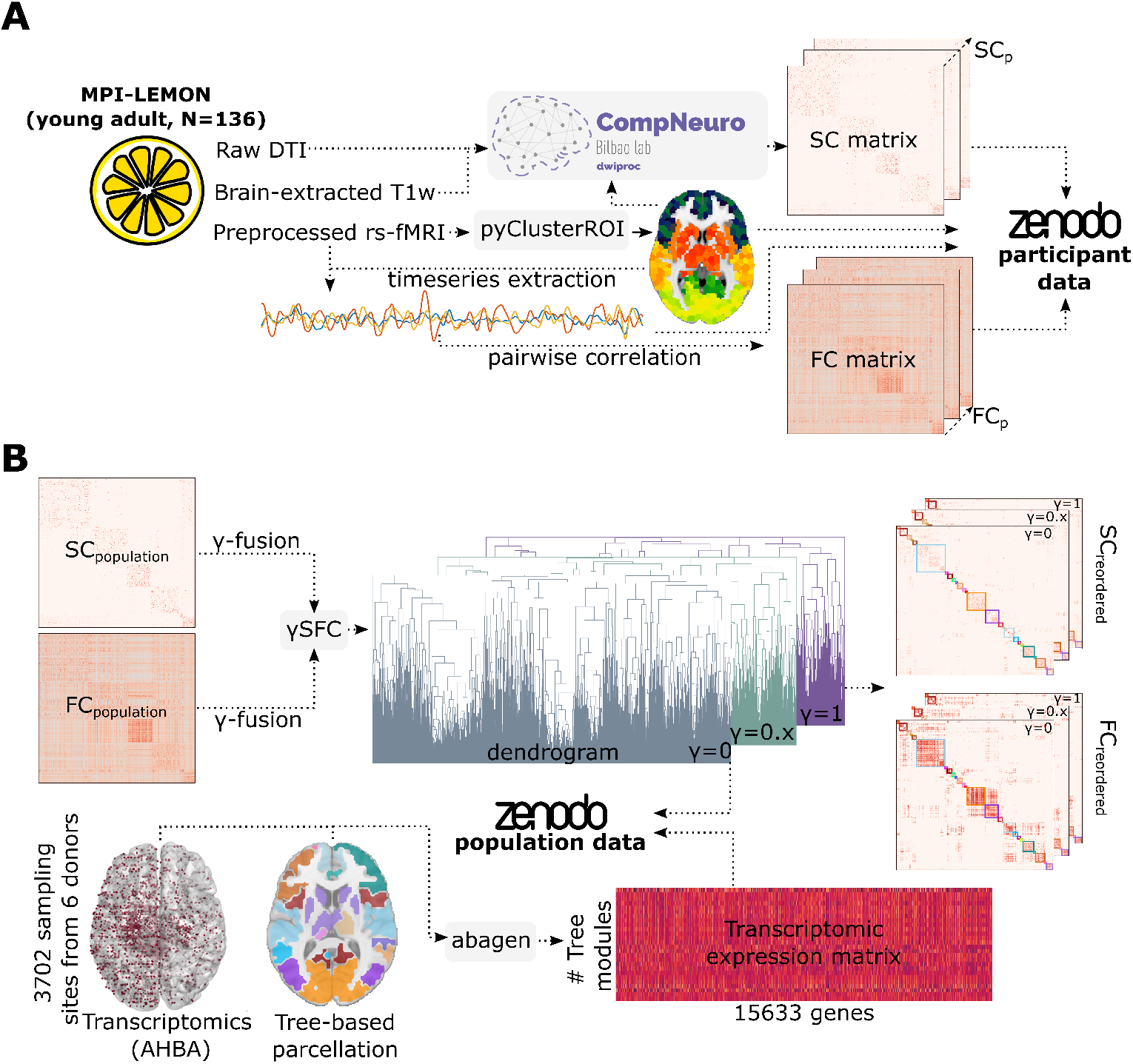
Methodological sketch and pipeline. **A:** Participants’ raw MRI data from N=136 healthy volunteers of the MPI-LEMON dataset were used. From preprocessed rs-fMRI images, we generated an initial brain parcellation using gray-matter masks and the open repository *pyClusterROI*. Subsequently, we employed our own publicly available code on GitHub to process the DWI images and extract the SC matrices. All post-processed data at the participant level, including the time series BOLD signal corresponding to the different iPAs, and the SC and FC matrices are available at https://zenodo.org/record/8158914. **B:** We calculated population SC_*p*_ and FC_*p*_ matrices, and subsequently generated *γ*-fused matrices as described in the methods section. The *γ*SFC matrices were used to construct dendrogram trees for different *γ* parameters. Additionally, we employed *abagen* to generate transcriptomic expression matrices for various tree-based parcellations. The transcriptomic matrices and dendrogram trees are also accessible on Figshare. The entire pipeline and project codes can be found on GitHub at https://github.com/compneurobilbao/bha2.

### Optimal γ-fused structure-function modular organization

The different trees of the *γ*SFC matrices describe distinct scales, enabling the construction of networks at different resolutions. The determination of the optimal partition by cutting the tree at M^*\**^ modules depends on the specific metric we seek to maximize. In order to showcase the versatility of our data with various metrics, we specifically concentrate on a metric we have previously defined and named cross-modularity^24^. Represented as *χ*, it is defined as the product of three quantities: the modularity of the functional matrix, the modularity of the structural matrix, and the average similarity between structural and functional modules.

Therefore, we aimed to maximize *χ* to select the optimal value of *γ* for a given initial parcellation atlas (iPA). Figure S1 shows box-plots of different values of the cross-modularity *χ* across different dendrogram levels and for different iPAs with different size, ie., varying the number of micro-regions ranging from 183 to 2165. For further analyses, we selected the iPA with 2165 micro-regions since it exhibited a higher mean value together with a lower variability value of *χ* within a wide range of scales, particularly within the first 120 levels where the maximum *χ* is located. Beyond this point, there was a decreasing trend. The optimal brain partition corresponds to the iPA that maximizes *χ* by reducing the dendrogram to M^*\**^ modules and setting the fusion parameter to *γ*^*\**^. In our case, the optimal value *γ*^*\**^ was determined to be 0.7, and this occurred at the level of 28 modules. However, two modules were considered invalid as they consisted of only one and two micro-regions, respectively, making it impractical to analyze the community structure within them. Henceforth, any subsequent analysis referring to the optimal brain partition will be based on the iPA that maximizes *χ* by reducing the dendrogram to M^*\**^ = 26 modules and setting the fusion parameter to *γ*^*\**^ = 0.7.

### Multi-scale γ-fused structure-function modular organization

As an illustrative example, and to gain a deeper understanding of how the multi-scale organization is influenced by *γ*, we investigated the network strength of the *γ*SFC matrix in two network representations. At the finest spatial resolution, corresponding to the lowest level in the hierarchical tree consisting of 2165 micro-regions (represented as brain maps in Figure 2A), and for each of the macro-regions, including the frontal lobe, parietal lobe, occipital lobe, temporal lobe, insula, and a collection of subcortical areas (Figure 2B). From both analyses, it is evident that increasing *γ*, transitioning from structure (*γ* = 0) to function (*γ* = 1), results in higher strengths shifting from subcortical areas to cortical regions. This pattern is observed consistently across all macro-regions, where the strength progressively increases with *γ*, peaking around *γ ≈* 0.7.

**Figure 2.**
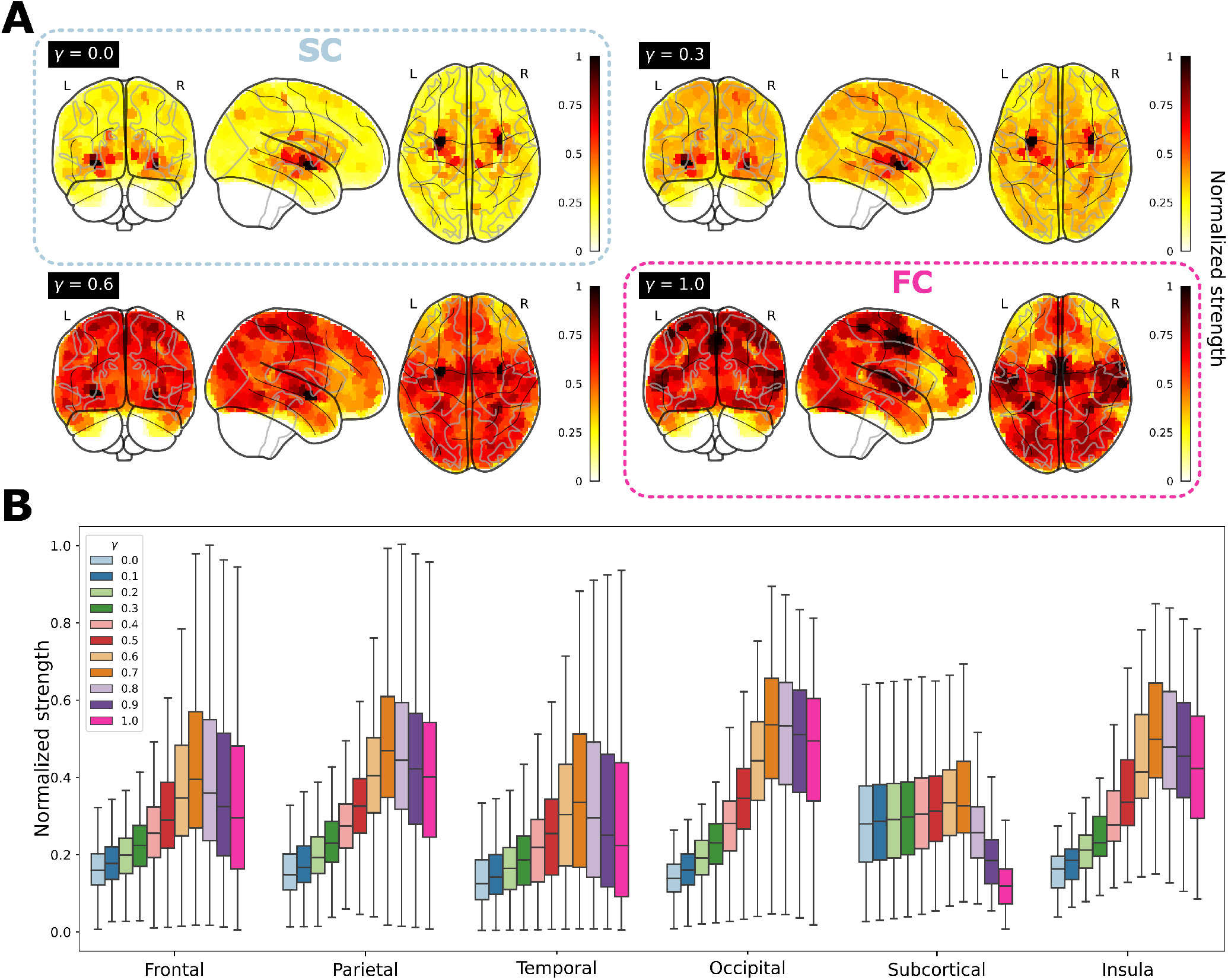
Graph-node strength is modulated by *γ*, the amount of interplay between structural connectivity (SC) and functional connectivity (FC). **A**: Brain maps of graph-node (normalized) strength. It is observed that for *γ* = 0 (pure SC), high strengths are predominantly localized in subcortical regions. However, as *γ* varies, there is a decrease in strength within these regions and an increase in cortical regions. **B**: Box plots for the distribution of graph-node strength across different macro-regions: Frontal lobe, Parietal lobe, Temporal lobe, Occipital lobe, a collection of Subcortical areas, and Insula. Notably, for each macro-region, the strength shows an increasing trend from *γ* = 0, reaching a peak at *γ ≈* 0.7, followed by a subsequent decrease.

Beyond the two selected levels of resolution for calculating strength, a hierarchical tree organization enables obtaining different metrics at different module-levels. For instance, we first colored the M^*\**^ = 26 distinct modules of the optimal brain partition for ease of visualization (Figure S2, top panel). Some tree metrics we computed were the module size (MS), height (H), and multi-scale index (MSI). While H is defined for each micro-region, MS and MSI are defined at the module-level. We also defined the module height (MH), by averaging over all individual H values within the given module (Figure S2, full dendrogram and dendrogram-inset). We represented these metrics on brain plots, where we observed distinct patterns for each module (Figure S2, brain plots).

We conducted further analyses to examine the correlations among these metrics and the intra-module strength, as a proxy for module segregation (Figure S3). From the different tree measures, both MS and MH showed a high correlation (r = 0.95, p < 0.001) with module segregation, as well as between themselves (r = -0.84, p < 0.001). Interestingly, the first module M1 was an outlier in the metric of MSI. Located at the border between the Precuneus, Isthmus Cingulate, and Posterior Cingulate cortices (Figure 3), M1 showed a remarkable resilience in remaining intact without splitting across multiple levels in the tree, possibly indicating a multi-scale functional role.

**Figure 3.**
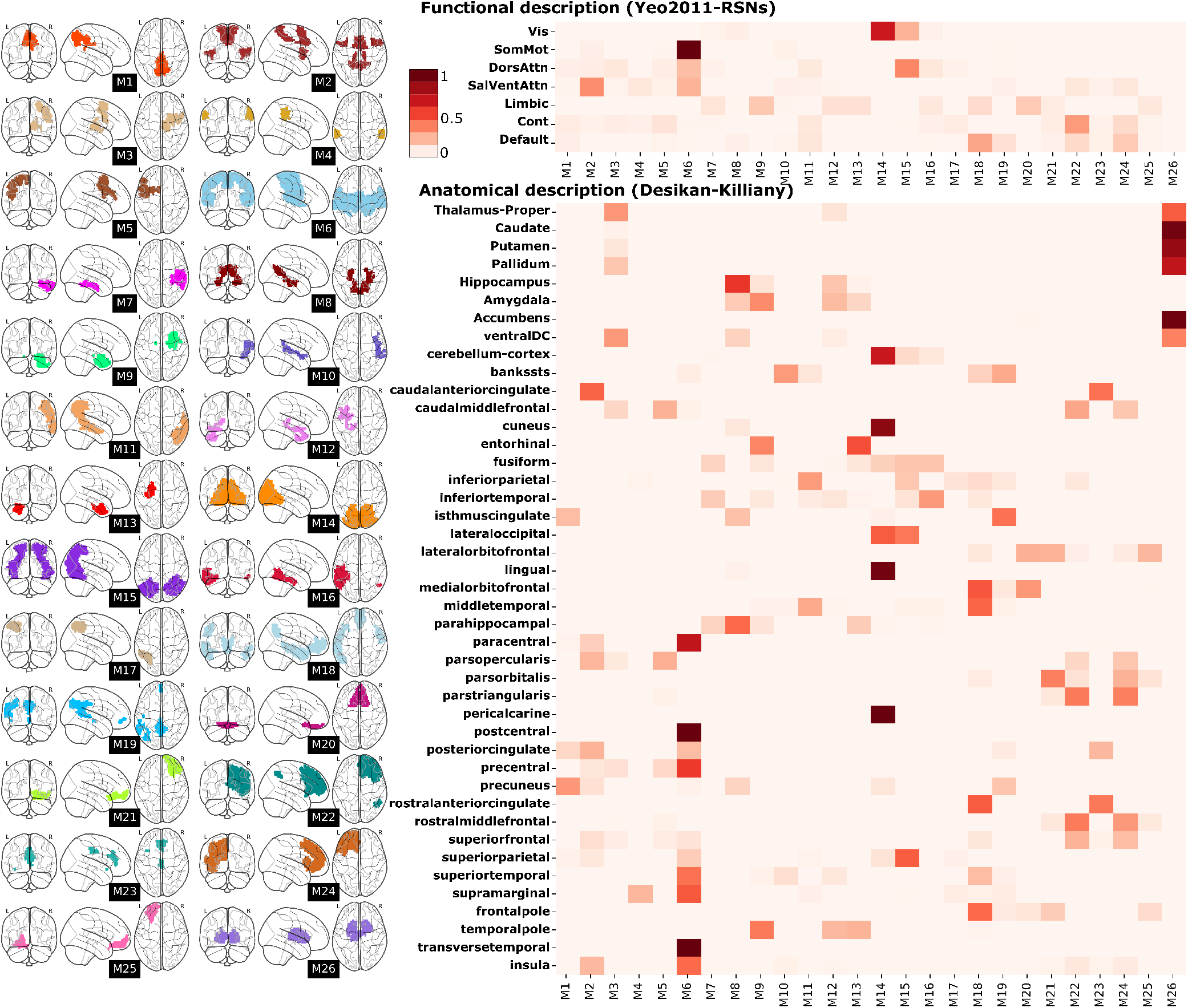
Brain localization, Functional description and Anatomical description of the optimal brain partition. **Left**: The different M1 to M26 modules belonging to the optimal brain partition are depicted within brain representations for localization purposes. **Right**: Additional information is provided regarding the specific location of each module, incorporating anatomical details and functional representations derived from the overlapping analysis among different modules and established Desikan-Killiany and Resting-State Network (RSN) atlases.

### Anatomical and functional description of a given brain partition

Our methodology for defining a given partition defined by specific values of M and *γ* is generally data-driven. At some point, it may be of interest to characterize the anatomical and functional properties of a specific partition. As an illustrative example, we made use of the optimal partition with M^*\**^ = 26 modules and *γ*^*\**^ = 0.7 and determined the spatial locations of these 26 modules within the brain (Figure 3, brain-glasses). We next obtained their functional and anatomical characterization by measuring respectively the amount of overlap within known Resting State Networks (RSNs)^59^ and regions from the Desikan-Killiany atlas^60^, the latter consisting of 34 cortical regions (in right and left hemispheres), 8 subcortical regions segmented from Freesurfer, and the cerebellum (Figure 3, heatmaps). Several of our modules exhibited strong overlapping within the RSNs. For instance, module M6 encompassed the Somato-motor Network (SMN) while also integrated parts of the Dorsal and Ventral Attention Networks (DAN, VAN). Module M18 showed an overlap with a portion of the Default Mode Network (DMN), and module M14 did it with the Visual Network (VIS). Moreover, certain modules displayed high overlapping with specific anatomical regions. For example, module M20 included the medial and lateral Orbitofrontal cortices, while module M26 integrated the Basal Ganglia and Thalamus.

### Neurobiological relevance of a given brain partition in brain-related disorders by making use of neurogenetics data

Similarly, it may be beneficial to characterize the modules of a given partition based on their participation in major brain-related disorders. To accomplish this, we focus again on the optimal partition and made use of publicly available transcriptomic data from the Allen Human Brain Atlas (AHBA)^47^, and constructed transcriptomic expression matrices at module-level (Figure 1B). In particular, we examined the transcriptomic expression within each module for genes associated with 32 different brain-related disorders, using a list of disorders and their corresponding genes extracted from the available data in^61^. For each disorder, the average expression across samples within a module was computed for the relevant genes, and z-score values were then calculated based on the optimal partition (Figure 4, heatmap). Modules with a z-score > 2, indicating high expression levels, or z-score < -2, indicating low expression levels, were marked with an asterisk. As the disorders were categorized into 7 disease groups (Psychiatric disorders, Substance abuse, Movement disorders, Neurodegenerative diseases, Tumor conditions, Developmental disorders, and Others), we also generated a z-score map representing the average expression of all genes associated with each disease group (Figure 4, brain-glasses). Modules M3 and M26 exhibited high relevance across multiple disorders. M26 represented a subcortical circuit encompassing the Basal Ganglia and Thalamus, while M3 represented a Thalamo-cortical circuit comprising a portion of the right Thalamus, a small segment of the right Putamen and Pallidum, and the right Caudal Middle Frontal, Superior Frontal, and Precentral gyri. M14 was relevant for 7 disorders and included the Occipital pole and the Medial Visual Cortex. Lastly, M8 displayed relevance in 6 disorders and represented a circuit connecting the Amygdala and Hippocampus with the Precuneus via the Parahippocampal and Isthmus Cingulate cortices.

**Figure 4.**
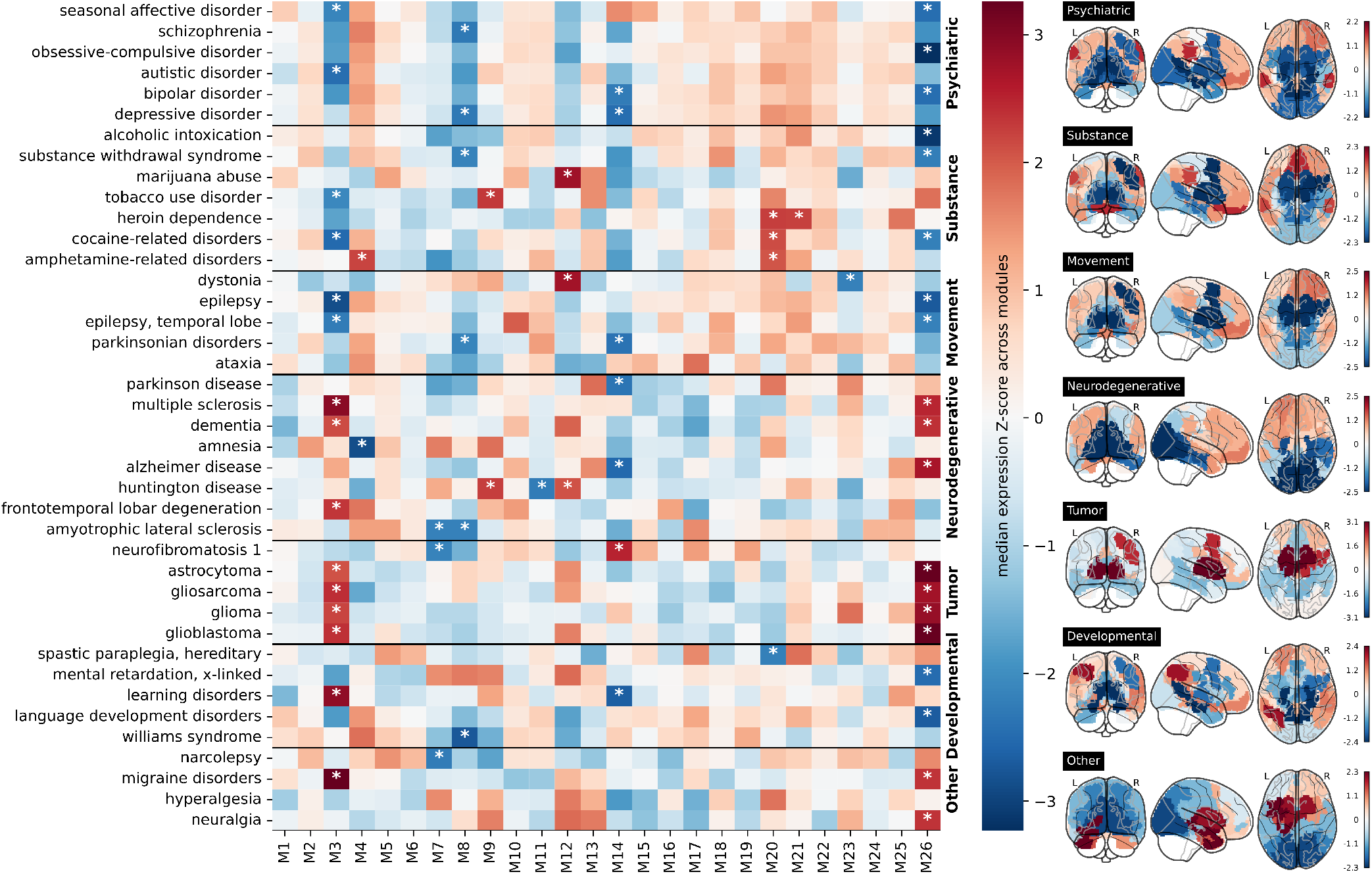
Neurogenetic module-level data for disease-related transcriptomic characterization of the optimal brain partition. **Left:** Within each module of the optimal brain partition (M1 to M26, x-axis), we depict the median transcriptomic expression values across different sets of genes associated with various brain-related disorders. Modules with significantly high/low expression values compared to the rest are denoted by a white asterisk (*). Notice that only specific modules among the 26 exhibited significant expression values. Notably, modules encompassing the basal ganglia and thalamus (M3 and M26) demonstrated significance across all disease groups. **Right:** Brain maps of median transcriptomic expression, averaged over different maps belonging to the same group of brain disorder. **Left, Right:** Z-scores were calculated across all the modules to indicate a relative high/low gene expression levels across modules.

## Discussion

In the words of Karl Friston, how rich functionality emerges from the invariant structural architecture of the brain remains a major mystery in neuroscience^7^. Previous research in network neuroscience has provided evidence for the emergence of hierarchical network modularity, facilitating the separation of neural computations at various spatial scales^13,18,21,26,31–34,36,38^. However, the specific role played by structure in functional organization, as well as the reciprocal influence, remain elusive. A key contributing factor to this gap in knowledge is the continuous interaction that takes place across multiple temporal and spatial scales^30^. Consequently, when employing a particular imaging modality for measurement, obtaining a comprehensive understanding of the intricate interplay between different scales becomes challenging.

For many years, our laboratory has been investigating the interplay between structure and function at the module-level. In our initial work 8 years ago^24^, we demonstrated excellent correspondence among modules, but not at the level of links in connectivity matrices. Since then, we have expanded our research to include studies on healthy brains^35,42,44^ and pathological brains^37,39–41,43,45,46^, exploring the multi-scale structural-functional correspondence at the module-level. Through collaborations with other groups, we have identified a lack of clear open data and repositories that facilitate this line of research. Many research groups in the field of physics and mathematics, specializing in complex network methodology, lack access to preprocessed brain networks derived from raw neuroimaging data. Motivated by this gap, our study aims to address this issue by employing state-of-the-art neuroimaging analyses, offering structural and functional networks at different scales, with both networks built using the same set of network nodes. Not only do we provide the final connectivity matrices, but also the code to generate them. Additionally, we introduce hybrid structure-functional networks through the parameterization of their overlap using the parameter *γ*. Furthermore, we have also released pre-processed neurogenetic data and code, to facilitate the connection between structural-functional correspondence across different scales and underlying biological mechanisms, opening up a promising avenue for studying brain disorders.

The structure of the released data, code and analyses are as follows:

- All preprocessed structural and functional connectivity matrices, transcriptomics data, and *γ*-dendrograms for the different iPAs have been uploaded to https://zenodo.org/record/8158914.
- The code to build *γ–*dendrogams is available at https://github.com/compneurobilbao/bha2/blob/main/src/build_tree.py.
- All notebooks needed to reproduce all figures of this manuscript are available at https://github.com/compneurobilbao/bha2/tree/main/notebooks

In this manuscript, which aims to be a practical guide for potential applications that make use of functional and structural connectivity matrices at different resolution scales, and their neurogenetic associations in relation to some brain pathologies, we have performed specific analyzes that have allowed us to produce 4 figures. Firstly, Figure 1 presents a methodological overview of the various datasets and key steps involved in our analyses. Figure 2 aims to demonstrate the continuous parametric effect of *γ* on different network metrics, and for that, we have depicted the distributions of node-graph strength at two different spatial resolutions. One resolution encompassed 2165 distinct microregions, while the other aggregated them into six distinct macro-regions covering the entire brain: Frontal lobe, Parietal lobe, Temporal lobe, Occipital lobe, a collection of Subcortical areas, and Insula. Figure 3 depicts a widely accepted strategy employed in neuroimaging studies for the comprehensive characterization of a specific brain partition, encompassing both functional and anatomical labels for each brain region. Finally, Figure 4 shows how gene transcriptomic imaging data can be used to characterize a given partition (in our example, the same partition chosen for Figure 3) in relation to a set of genes with significant associations in 32 different brain disorders. These disorders are grouped into various categories such as psychiatric disorders, substance abuse, movement disorders, neurodegenerative diseases, tumor conditions, or developmental disorders.

To summarize, with this study we aim to provide valuable resources in the form of datasets and code to facilitate the exploration of the intricate interplay between brain structure and function across diverse spatial scales, including the aggregated level. It is our intention to contribute to unraveling the enigmatic nature of this structure-function relationship, which, despite significant progress, still harbors substantial gaps in our understanding. We encourage other researchers with an interest in this area to make use of the resources presented here to further contribute to the elucidation of this fascinating phenomenon.

## Material and methods

### Raw neuroimaging dataset

The Max Planck Institut Leipzig Mind-Brain-Body Dataset, commonly referred to as LEMON^58^, is a comprehensive multimodal dataset encompassing MRI sequences, EEG, ECG, and behavioral scores. In this study, we focused on a subgroup of N=136 healthy individuals within the age range of 20 to 30 years, with a male population comprising 98 individuals (72%). For the precise selection of the IDs individuals, see https://zenodo.org/record/8158914/files/participants.tsv. Specifically, for this study we made use of three different MRI sequences: T1, rs-fMRI, and DWI, with the aim of extracting functional and structural connectivity matrices. For a comprehensive description of the sequences employed, please refer to Table 7 in^58^.

### Preprocessed neuroimaging dataset

LEMON also provides access to preprocessed data. In our study, we made use of the brain-extracted T1 and rs-fMRI images as input data for further analysis. For preprocessing DWI images, we made used of a custom pipeline utilizing the the brain-extracted T1 images, MRtrix3 (*v*3.0_*RC*3), FSL (v6.0.1), and ANTs (v2.3.1). Our DWI pipeline involved cleaning the raw images using *DWIdenoise* and *DWIpreproc* tools, which included correction for eddy current distortion and susceptibility-induced distortion using the *topup* tool. Next, we performed whole-brain probabilistic tractography using the *iFOD2* tracking algorithm in MRtrix3. The tractography was initiated from seeds located at the gray matter-white matter interface, with a selection of 3 million fibers based on an angle threshold of < 45º and a length threshold of < 200 mm, following the recommendations in the MRtrix3 documentation.

### Data-driven anatomical and functional solutions for different initial brain partitions

Following a methodology similar to^24^, and utilizing the preprocessed rs-fMRI images, we performed an unsupervised voxel-level clustering analysis to delineate a specific number of micro-regions following the strategy proposed in^62^. These micro-regions define the highest spatial resolution scale employed for subsequent analyses. In contrast to^24^, where all brain voxels were included into the same clustering process, our methodology here involved six separate clustering procedures, each one focused solely on the voxels contained within distinct macro-regions, namely the frontal lobe, parietal lobe, occipital lobe, temporal lobe, insula, and subcortical structures (pooling together the thalamus, basal ganglia, amygdala, and hippocampus). Thus, while a functional partition was applied, the resulting regions were anatomically constrained by the defined macro-regions. By aggregating all micro-regions obtained from the six different macro-regions, we generated an initial Parcellation Atlas (iPA). We followed the same procedure to generate 9 different iPAs, each consisting of a different number of micro-regions, specifically 183, 391, 568, 729, 964, 1242, 1584, 1795, and 2165.

### Brain functional and structural connectivity matrices, and their fusion through the parameter γ

FC matrices were generated by computing the Pearson correlation between pairs of regions fMRI time-series, each one obtained by averaging voxel-wise fMRI time-series within each micro-region within a specific iPA. For SC matrices, we employed the *tck2connectome* tool to perform fiber counting of the streamlines connecting pairs of micro-regions in the iPA. Prior to constructing SC, a nonlinear registration of the iPA to the b0 image of the DWI sequence was performed.

From the individual SC and FC matrices, we computed population matrices SC_*p*_ and FC_*p*_ by calculating for each link the median value across all equivalent links in the individual matrices. Next, and to match the sparse nature of SC_*p*_, a threshold was applied to FC_*p*_, resulting in comparable link density. Subsequently, we binarized the two matrices and fused them into a single matrix as follows:

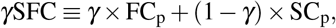

where *γ* is a real number between 0 and 1. This strategy allows us to continuously parameterize the level of interplay between SC_*p*_ and FC_*p*_, enabling the recovery of each connectivity class when *γ* = 0 or *γ* = 1, while obtaining hybrid connectivities for intermediate values.

### Multi-scale hierarchical representation at module-level

For each of the 9 distinct iPAs, a hierarchical agglomerative clustering (HAC) was employed to extract different nested modules, allowing to build different network resolutions at different scales (each one determined by a cutting tree number of modules M). More specifically, HAC was performed using a weighted method on the connectivity patterns obtained from the matrix *γ*SFC, here defined as the correlation distance between pairs of micro-regions in the *γ*SFC matrix.

Within the context of tree or dendrogram partitions, different multi-scale metrics can be established. These metrics can be defined either at the module-level or at a more granular level (with the highest spatial resolution determined by the micro-region level). In this particular study, we have defined the module size (MS), representing the count of micro-regions encompassed within a module. Additionally, we have introduced the multi-scale index (MSI), which quantifies the number of levels within the tree structure where a module remains intact without further division. In relation to micro-regions, we have selected the height (H) metric, which denotes the specific level at which a given micro-region separates from its parent module.

### The optimal brain partition based on cross-modularity maximization

After obtaining a hierarchical partitioning of the connectivity matrix, clustering micro-regions into distinct modules M, the determination of the optimal level for cutting the tree depends on the specific metric being optimized. In this study, following the methodology introduced in^24^, the metric chosen for maximization is the cross-modularity *χ*, defined as:

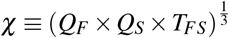

This metric simultaneously accounts for three different qualities: the modularity of the functional matrix (*Q*_*F*_), the modularity of the structural matrix (*Q*_*S*_), and their similarity (*T*_*FS*_). The latter is defined as the DICE similarity between a functional module and a structural module, and averaging across all modules in the given partition. By varying M along the tree, we can calculate *χ* for each configuration M and select the optimal partition M^*\**^ where *χ* is maximized. Additionally, unlike the approach in^24^, we here have introduced a second parameter *γ*^*\**^ in the maximization of *χ*, which controls the amount of structure-function interplay.

### Transcriptomic data at module-level to assess brain-related disorders

In addition to multi-scale brain partitions and structure-function analysis, we processed the transcriptomic open data from the Allen Human Brain Atlas (AHBA)^47^ by using the *abagen* tool developed in^54^. In particular, *abagen* allows us the generation of brain transcriptome module-level values for a specific parcellation, that in our case, we chose the optimal brain partition. Moreover, we examined the transcriptomic expression at module-level in relation to genes associated with 32 brain disorders introduced in^61^. To do this, we calculated the mean transcriptomic expression of all genes related to each disorder within each module. Subsequently, we performed a z-score analysis for each disorder to identify modules with low transcriptomic expression (z < -2) and modules with high transcriptomic expression (z > 2). The average expression of the different genes were also grouped into 7 different disease cateogories: Psychiatric disorders, Substance abuse, Movement disorders, Neurodegenerative diseases, Tumor conditions, or Developmental disorders.

## Data Availability

This work involves data from different sources. First, the raw MRI images can be downloaded from the LEMON database webpage http://fcon_1000.projects.nitrc.org/indi/retro/MPI_LEMON.html. Second, the transcriptomic AHBA expression data can be downloaded from https://human.brain-map.org/. All post-processed data obtained in this manuscript, including the time-series fMRI signals corresponding to regions in the different iPAs, the SC and FC matrices, the optimal brain partition, and the *γ*-modulated trees are available in https://zenodo.org/record/8158914.

## Code Availability

We have used open tools available on GitHub, and we have also uploaded our own tools and code used in this manuscript. In particular,

- For defining the iPAs, we have used the open tool “pyClusterROI” (https://ccraddock.github.io/cluster_roi/).
- For obtaining the transcriptomic expression matrices, we have used *abagen* (https://abagen.readthedocs.io/en/stable/).
- For preprocessing the DWI images and obtaining the SC matrices, we have used our own code (https://github.com/compneurobilbao/compneuro-dwiproc).
- All the analyses and code involved in the methods and results of this manuscript have been uploaded to https://github.com/compneurobilbao/bha2.

## Acknowledgements

This work would not have been possible without the tremendous effort of numerous colleagues with whom we have actively contributed over the past 8 years to various studies exploring the multi-scale nature vs. optimal partitions of structure-function multi-scale networks. The list of researchers includes: N. Aginako, C. Alonso-Montes, J.C. Arango-Lasprilla, P. beim Graben, M.P. Boisgontier, B. Camino-Pontes, Hu. Chen, He. Chen, R. Cofré, M. Desroches, D. Drijkoningen, X. Duan, F.E. Rosas, I. Escudero, M. Fernandez, I. Fernandez-Iriondo, I. Gabilondo, M. Gatica, J. Gooijers, X. Guo, C. He, X. Huang, L. Li, W. Liao, D. Marinazzo, B. Mateos, P.A.M. Mediano, M.A. Muñoz, L. Olabarrieta-Landa, P. Orio, S.P. Swinnen, L. Pauwels, J. Rasero, L. Remaki, S. Rodrigues, F.E. Rosas, W. Sheng, B. Sierra, L.Q. Uddin and J. Xiao. AJM is funded by a predoctoral contract from the Department of Education of the Basque Country (PRE-2019-1-0070). JMC is funded by Ikerbasque: The Basque Foundation for Science, and from the Health Department of the Basque Country (grant 2022111031).

## Author contributions statement

J.M.C., P.B., I.D., A.E., S.S. and A.J.M. designed the project. A.J.M. performed the data analyses. S.S., P.B., and J.M.C. supervised the methodology. A.J.M. and J.M.C. made the figures. A.J.M. and J.M.C. drafted the manuscript. All authors wrote and reviewed the manuscript.

## Supplementary Figures

**Figure S1.**
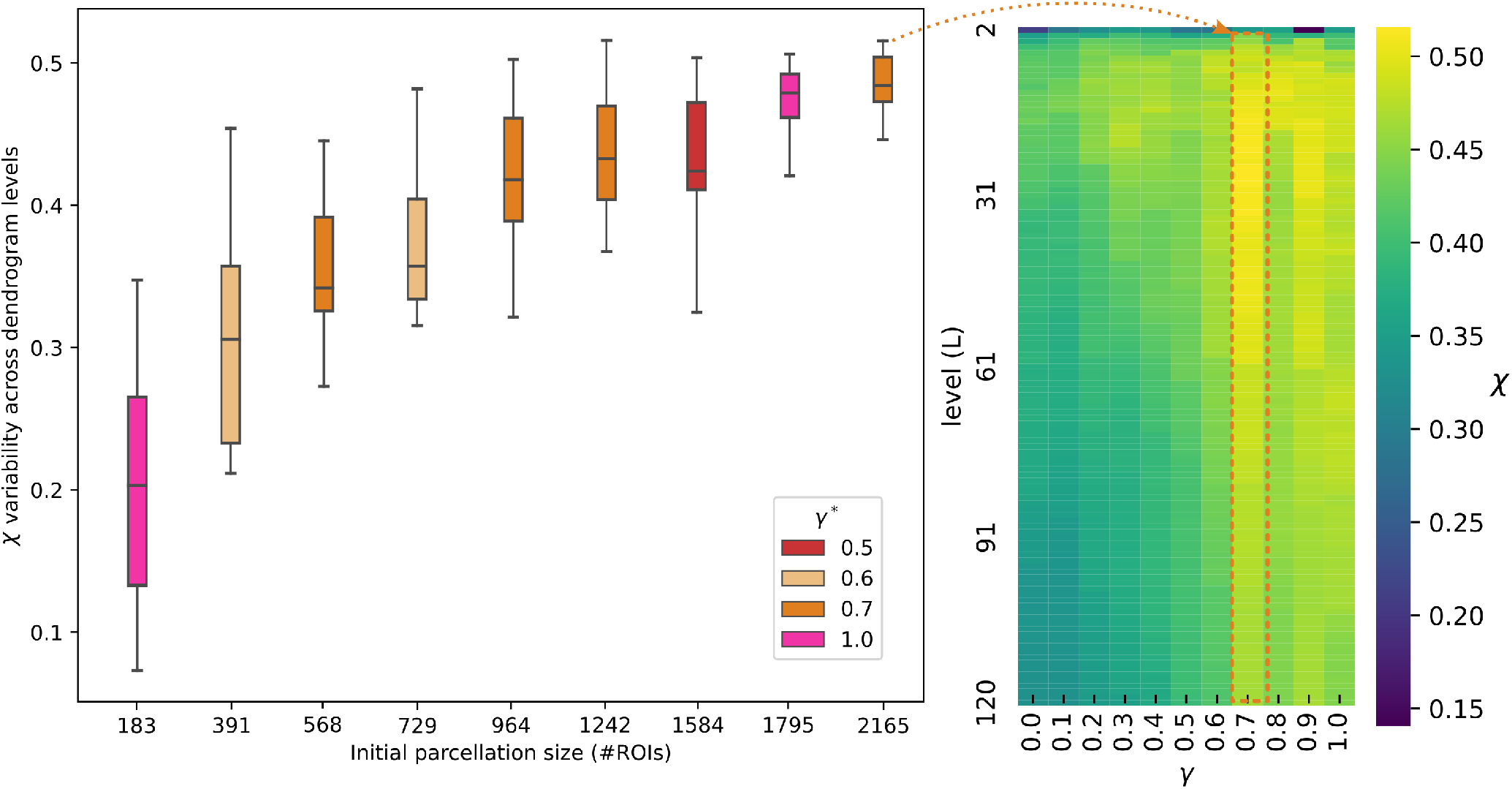
Selection of best initial parcellation atlas and optimal value of *γ* parameter. **Left:** Box-plots of the cross-modularity *χ* across different dendrogram levels and for different size of the initial parcellation atlas (iPA). The different values of *χ*, on which the box-plots are calculated, come from the different levels in the dendrogram (here we have varied from 2 to 120). As the number of ROIs for the initial parcellation increases, the cross-modularity *χ* also increases. **Right:** The optimal value of *γ*^*\**^ = 0.7 is also illustrated for the same range of dendrogram levels, ie. from 2 to 120.

**Figure S2.**
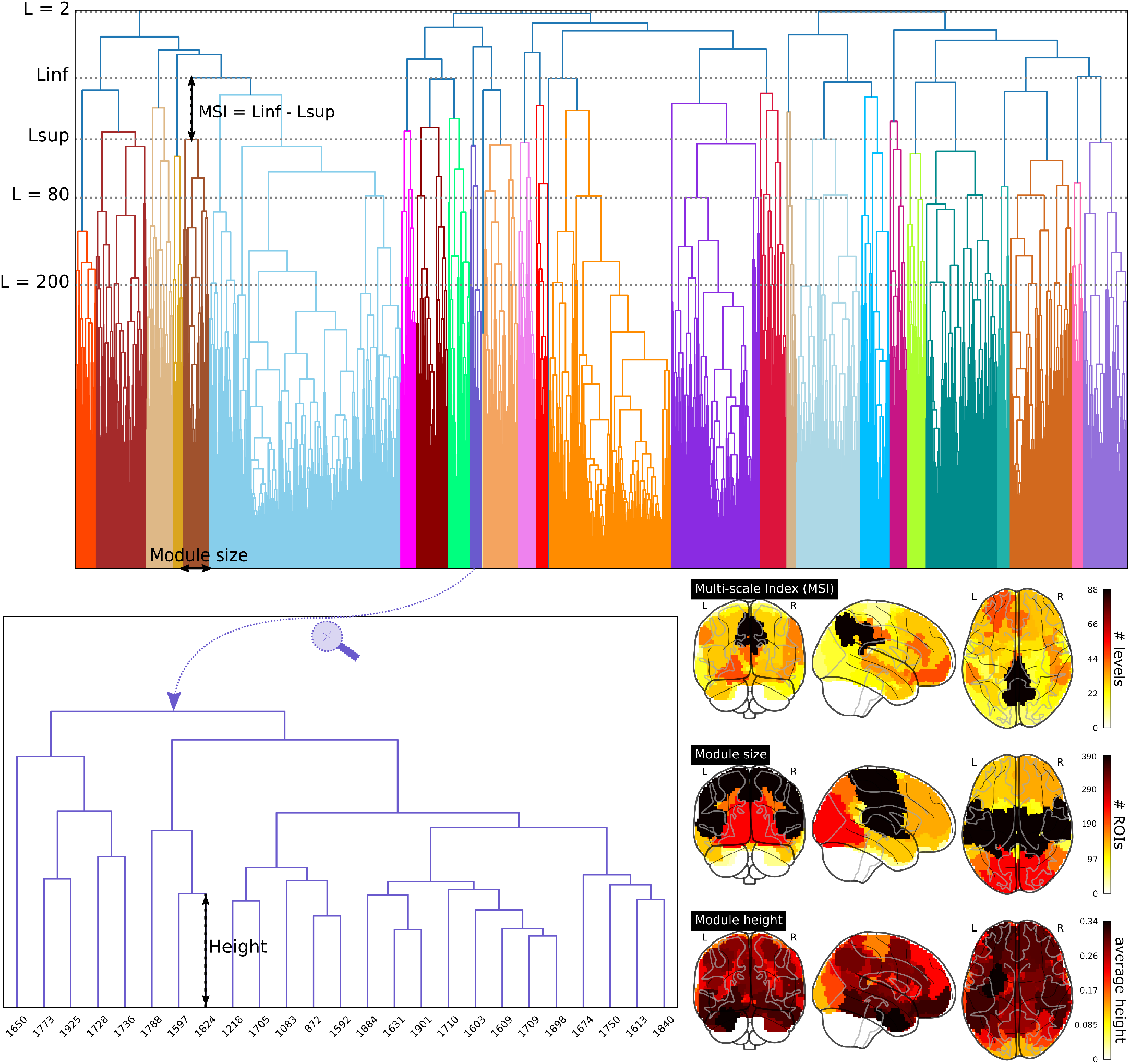
Dendrogram measures and module analysis in the the optimal brain parcellation. Each color in the dendrogram corresponds to a distinct module (that for the optimal brain partitions correspond to 26 different modules). The y-axis of the dendrogram illustrates different levels (L) and provides visual representation of a given module and their inferior (Linf) and superior (Lsup) levels. Different multi-scale metrics can be defined, eg., higher values of the Multi-scale Index (MSI) correspond to enhanced preservation of the module throughout multiple levels in the dendrogram, indicating heightened stability across the tree structure. The module size is also represented by the width of each module in the dendrogram. The lower inset shows a marked micro-region Height indicating to which extent a given micro-region is detached from the overall tree structure. Lower values of Height represent stronger connectivity of the micro-region with the other micro-regions belonging to the same module. The figure also shows brain maps depicting the three aforementioned measures for each of the modules.

**Figure S3.**
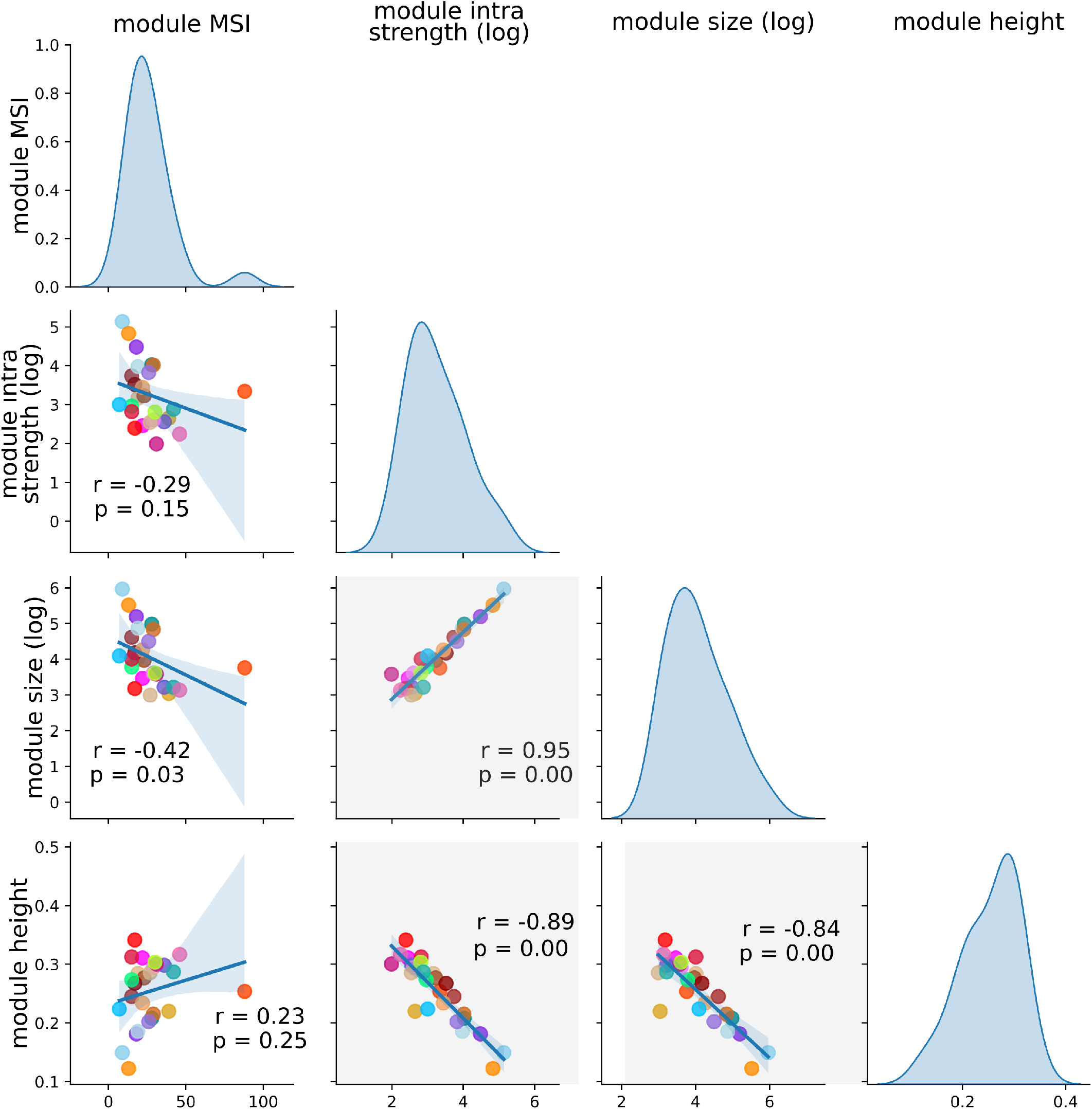
Statistical dependencies between different multi-scale metrics. Cross-correlation plots between multi-scale index (MSI), intra-strength (as a proxy for module segregation), size and height. Different color-points represent different modules of the optimal brain parcellation with a total number of 26 modules. Principal-diagonal plots show probability density functions of the different metrics. Notice that higher correlations between module size, module height, and intra-strength indicate that heightened connectivity within modules (segregation) results in larger dendrogram modules and delayed micro-region splitting in the tree. The MSI reveals a distinct outlier (M1) encompassing several structures such as a part of the precuneus, isthmus cingulate, and posterior cingulate, located at the border between DMN, fronto-parietal, and dorsal attention networks.

